# Lateral Flow Assay Sensitivity and Signal Enhancement via Laser µ-Machined Constrains in Nitrocellulose Membrane

**DOI:** 10.1101/2024.05.09.593095

**Authors:** Gazy Khatmi, Tomas Klinavičius, Martynas Simanavičius, Laimis Silimavičius, Asta Tamulevičienė, Agnė Rimkutė, Indrė Kučinskaitė-Kodzė, Gintautas Gylys, Tomas Tamulevičius

**Affiliations:** Department of Physics, Kaunas University of Technology, Lithuania; Institute of Materials Science, Kaunas University of Technology, Lithuania; UAB Sanpharm, Lithuania; Institute of Biotechnology, Life Sciences Center, Vilnius University, Lithuania

**Keywords:** lateral flow assay, nitrocellulose, μ-channels, laser µ-machining, reaction time, calorimetric sensing, signal enhancement, SARS-CoV-2

## Abstract

Multiplex lateral flow assay (LFA) is a handful diagnostic technology that can identify severe acute respiratory syndrome coronavirus 2 (SARS-CoV-2) and other common respiratory viruses in one strip, which can be tested at the point-of-care without the need for equipment or skilled personnel outside the laboratory. Although its simplicity and practicality make it an appealing solution, it remains a grand challenge to substantially enhance the colorimetric LFA sensitivity. The local flow rate constraints imposed in nitrocellulose (NC) membranes via a number of vertical femtosecond laser micromachined microchannels are important for prolonged specific binding interactions. Porous NC membrane surfaces were structured with different widths and densities μ-channels employing a second harmonic of the Yb:KGW femtosecond laser and sample XYZ translation over a microscope objective-focused laser beam. The influence of the microchannel parameters on the vertical wicking speed was evaluated from the video recordings. The obtained results indicated that μ-channel length, width, and density in NC membranes controllably increased the immunological reaction time between the analyte and the labeled antibody by 950%. Image analysis of the colorimetric indicators confirmed that the flow rate delaying strategy enhanced the signal sensitives by 40% compared with pristine NC LFA.

**Graphical abstract:** 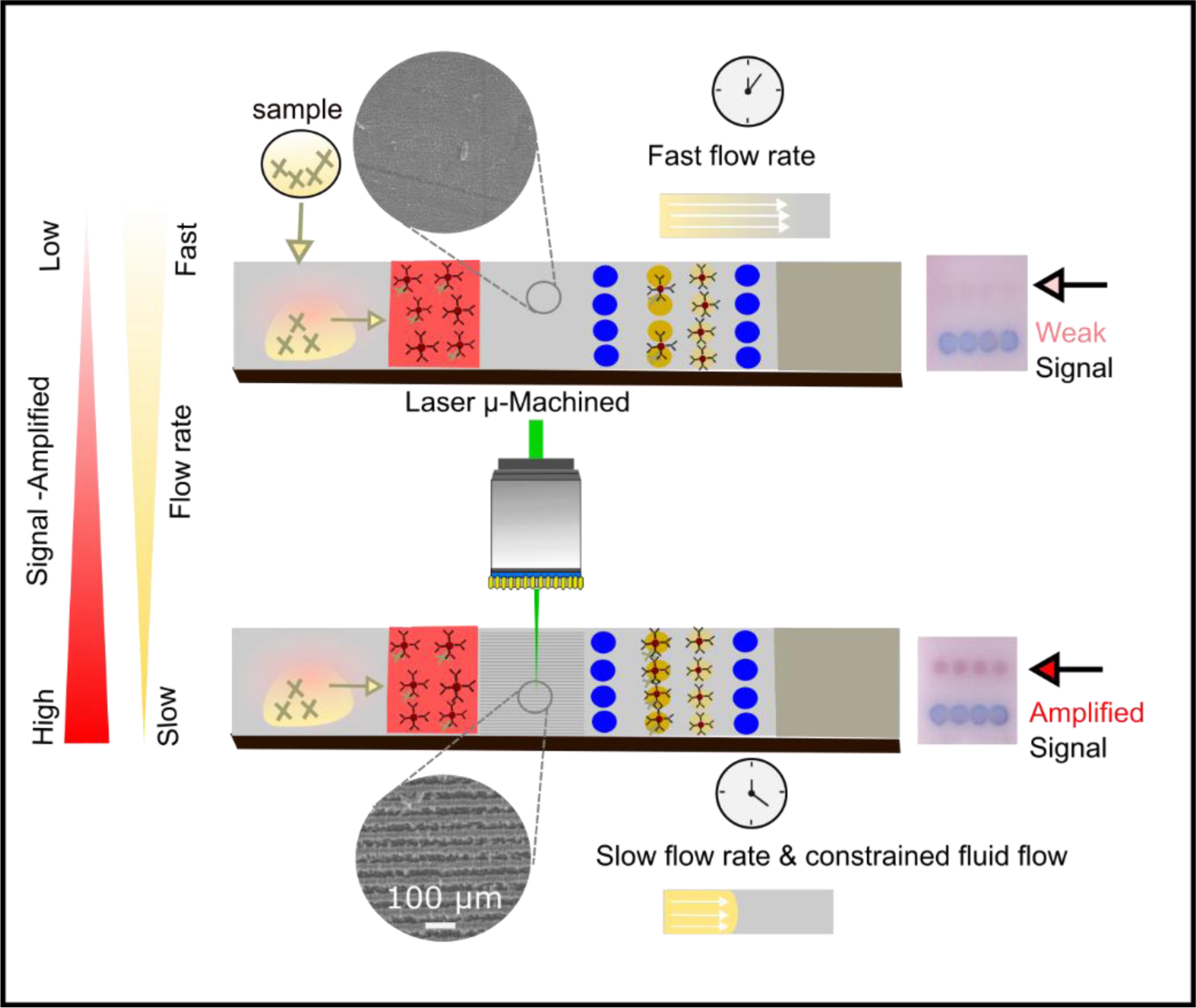

## 1. Introduction

On the verge of viral disease outbreaks effective diagnostic methods are of paramount importance. Point-of-care (PoC) diagnostic instruments as rapid diagnostic tests are expected to possess the following attributes: affordability, sensitivity, specificity, user-friendliness, rapidity, robustness, equipment independence, and accessibility for end-users which was abbreviated as ASSURED criteria by the World Health Organization (WHO) in 2003 [1]. Recent progress and availability of smart devices introduced the possibility for real-time connectivity (R) together with ease of specimen collection and environmental friendliness (E) translating them into REASSURED acronym[2]. Even though many accurate, reliable, and highly sensitive laboratory methods can be used at the single-cell level such as polymerase chain reaction (PCR), enzyme-linked immunosorbent assay (ELISA), flow cytometry, and Raman spectroscopy, these methods do not fully meet REASSURED criteria. Over nearly the past two decades, the PoC diagnostic methods have made significant progress toward meeting the outlined criteria. In particular, lateral flow assays (LFAs) have emerged as a prominent solution due to their simplicity, speed, portability, affordability, a wide variety of applications, shelf stability, and ease of use [3].

Currently, LFAs serve as the most common PoC sensors for diagnosing viruses (human immunodeficiency virus (HIV), hepatitis, severe acute respiratory syndrome coronavirus 2 (SARS-CoV-2) [3]), bacteria (*streptococcus*, *E. coli*), and parasites (malaria [4]) caused infections. They are widely used for determining medical conditions (pregnancy testing and cardiac markers), environmental monitoring (pesticides and heavy metals [5]), food safety applications (such as allergens and pathogens [6]), and cancer markers [7] (prostate-specific antigens, alpha-fetoproteins [8]). The LFAs are popular because they are cheap and simple but at the same relatively accurate PoC method. The backbone of LFAs is a nitrocellulose (NC) paper-based substrate that enables low-cost and sustainable manufacturing. The NC membrane is a microporous material that allows for capillary forces to drive the fluid flow through the test strips where the analytes are detected via specific reactions. The natural capillary action in NC eliminates the need for external pumps or complex microfluidic systems. Porous NC matrix provides equipment-free passive microfluidics at the same time providing a biocompatible scaffold that is well-suited for facilitating essential antibodies and antigens biointeractions. Additionally, the flexibility of paper substrates allows for the development of diverse nano-biosensing designs and strategies, *e.g.*, barriers, and constrictions [9].

LFA traditionally consists of a sample pad, NC membrane, conjugate pad, absorbent pad, and polyvinyl chloride backing card (see suplementarty **Figure S1**). Commercially, the whole assembly is embedded into a housing chamber to improve its usability. The sample and adsorbent pads are primarily made from cellulose or glass fiber, and the NC is the most relevant material for the reaction zone. Glass fiber is frequently employed for conjugate pad preparation. The fundamental principles and methodologies of traditional LFAs have been extensively discussed in various scholarly articles and training manuals and will not be discussed in more detail [9].

Although LFAs exhibit significant versatility, their use is constrained to mainly qualitative analysis and detection of analytes with relatively high 0.1 mg/mL concentrations. The LFAs are not optimally designed for quantitative measurements or the analyte presence identification at extremely low concentrations as 0.1 ng/mL or fg/mL. The limited ability to identify low-concentration analytes is a significant limitation of LFAs [10].

To overcome these constraints, numerous strategies have been employed to enhance the sensitivity and specificity of LFAs to achieve more precise and superior PoC diagnostics. The most recent research in LFAs concentrates on improving the limit of detection (LoD) and sensitivity [11] as well as multiplexing capabilities for detecting multiple analytes from a single sample [12]. The first group of strategies is related to nanoparticle (NP) development. Red-appearing gold NPs are traditional colorimetric labels used for LFAs. The visual appearance of noble metal NPs depends on size, shape, elemental composition, exterior structure, interparticle distance, and dielectric constant as it is related to the localized surface plasmon resonance (LSPR) condition. For example, increasing the size of the NPs results in a more sensitive colorimetric signal output [9]. In addition, when compared to spherical NPs of similar size, anisotropic NPs with more intricate structures alleviate visual readout of the plasmonic signals and improve the LFAs detection sensitivity. Alloyed NPs including Au@Pt [13], Pt@Pd [14], and Pt@Ni [15], have enhanced the analytical capabilities of LFAs. For example, Guodong Liu *et al.* created Au-Pt nanoflowers serving as colored and catalytic markers for highly sensitive microRNA-21 assay LFA applications where 0.3 pM LoD was achieved in under 15 minutes [13].

The second effective LoD enhancement strategy was the sample enrichment method where the LFA strip construction was modified by placing the conjugate pads after sampling, as opposed to traditional LFAs [11, 16]. It resulted in a significantly larger concentration of the probe-analyte conjugate in the test zone as a result of the elevated capture rate of Au NPs compared to the direct sampling method. The LoD for microRNAs was enhanced by a factor of 10 to 100 compared to the standard LFAs [17]. The third LFA performance-enhancing strategy that is closest to the one discussed in this work is local control of the lateral flow speed [11]. It influences the reaction-specific kinetics resulting in the color intensity changes at the detection site. Flow rate can affect reagent dissolution and mixing, as well as reaction effectiveness, and ultimately impact the specific binding (SB) and nonspecific binding (NSB) events, which, in turn, determine the sensitivity and specificity of the assay. Understanding the influence of lateral flow speed on the reaction kinetics and binding events provides a foundation for exploring how reactants such as analytes, labeled detection particles (like Au NPs), and various buffer components travel through the NC membrane. In LFAs, specific binding interactions are targeted to occur at designated test and control lines. This transport within the membrane is governed by mass transfer mechanisms including advection, diffusion, and dispersion.

Advection is the primary mechanism for the initial movement of reactants from the sample application point toward the test and control lines. It involves the transport of substances by the bulk movement of a fluid - usually a liquid sample like blood, serum, or saliva - that carries the reactants through the membrane via capillary action. The directional flow, influenced by the local NC properties, such as flow rate adjustments, significantly affects how and where reactants move within the assay setup. Diffusion, although slower than advection, plays a crucial role in distributing reactants within the membrane, particularly across its width. This ensures that reactants evenly interact with the receptors on the test line, crucial for achieving consistent assay results.

Lastly, dispersion, which can be seen as a combination of advection and diffusion, occurs due to variations in flow velocity within the porous structure of the NC membrane. This effect leads to a broadening of the reactant front, enhancing interaction with line receptors but potentially decreasing the sharpness of the detection lines. Understanding these mechanisms helps in refining LFA design and execution, ultimately improving diagnostic accuracy and reliability [10].

The kinetics of transport and immunological reaction are characterized by the Péclet number (Pe) and the Damköhler number (Da), respectively [9]. Where The Péclet number is a dimensionless number that characterizes the relative importance of advective (or convective) transport to diffusive transport. The Damköhler number is a dimensionless number that compares the chemical reaction rate to the rate of transport processes (such as diffusion). In LFAs, a high *Pe* (≫ 1) indicates that advection (fluid flow) significantly contributes to the Au NPs transport towards the test site, compared to diffusion. This is important for ensuring that the Au NPs can reach the test site in a timely manner. A low *Da* (≪ 1) indicates that the reaction rate (*e.g.*, the binding of Au NPs to target molecules at the test site) is the limiting step, not the transport of Au NPs to the site. This could imply that enhancing the reaction kinetics, for example, by increasing the affinity of the Au NPs for the target or optimizing the test site conditions, could improve the assay’s performance [18].

The advective flux depends on the velocity of the fluids flowing inside the LFA membrane (*v*), the pore size of the NC membrane (*S*), and the diffusivity of the molecules (*D*) (see **Table S1** for details). The reaction rate depends on the concentration of capture molecules in the test region (*C_e_*), the forward reaction rate constant of the single antibody-antigen immunoreaction (*K_on_*). Typical parameter values from the literature are summarized in **Table S2**.

LFAs are intrinsically limited by the reaction rate, and, thus, improving the reaction efficiency is the most critical step toward maximizing SB and boosting the sensitivity of LFAs. The diffusion rate limits the transport of molecules and labels, and the surface reaction rate limits the surface reaction.

Increasing reaction efficiency can be accomplished by increasing the reaction rate where the slowing down of the lateral flow increases the reactant concentration and prolongs the reaction time [11].

Among the most important reasons for the poorer sensitivity of LFAs in comparison to other standard laboratory tests, such as ELISA, is that they do not provide as much time for specific binding of biorecognition receptors to antigens or target markers. This is due to the inherent capillary effect of NC (also known as wicking), which creates a unidirectional flow of the liquid solution at a constant value across the membrane [16]. The investigated reaction kinetics are associated with forming the conjugation/antigen/captured antibody ternary (sandwich ternary). Liang *et al.* observed that sandwich ternaries form more slowly when a conjugated label is first used to bind the antigen and then a capture antibody is added sequentially [19]. Thus, the LoD for malarial protein on a sequential flow was reported to be 4 to 10 times lower than that on a premixed flow [19]. Therefore, the goals of optimizing the assay kinetics are to (i) maximize SB and (ii) minimize NSB. Quantitative assay optimization can be achieved by maximizing the signal-to-noise ratio (SB/NSB) [11].

Several studies have been conducted on improving reaction kinetics. At least two methods were reported for reducing the flow rate in LFAs: modifying NC chemically and changing the geometry of the components. Feng Xu *et al.* slowed down the flow by incorporating a salt barrier. It was found that adding a saline barrier to the membrane before the test region expedited the hybridization reaction attaining a tenfold enhancement sensitivity but at the expense of reduced flow velocity and an extended assay time [20]. Cellulose nanofibers can also be used to modify membranes by changing the properties of the NC membrane to enhance the biomolecule adsorption ability boosting detection sensitivity by

20-fold [21]. Cellulose nanofibers were also reported to bring the captured molecules closer to the surface, thus increasing the colorimetric intensity of the captured labels by 36.5% [22]. Yew *et al.* utilized electrospinning to deposit polycaprolactone (PCL) nanofibers creating a hydrophobic region on NC membrane and decreasing the flow rate from 0.35 mm/s to 0.29 mm/s [23]. Alternatively, Rivas *et al.* implemented wax pillars on the NC surface to create a delay in the flow and induce pseudo turbulences in the capillary flow. Their approach led to a 3-fold increase in sensitivity for HIgG detection, compared to the traditional LFA method [24].

In contrast, Katis *et al.* have devised a method to decrease the width of the Test line from 5 to 1 mm. A laser was employed to induce the polymerization of a specific pattern on the NC membrane. By constraining the flow route, a simultaneous augmentation in both the duration of flow and the concentration of the reagents was achieved resulting in a significant 30-fold increase in sensitivity for detecting C-reactive protein. The laser micromachining process is cost-effective and is favorable when compared to alternative patterning techniques like photolithography [25].

To summarize, the process of modifying assay kinetics is a critical step in improving LFAs, which has the potential to increase sensitivity by several orders of magnitude. Flow speed control enables reaction rate control, ensures reactant concentration, prolongs the overall reaction duration, and augments the number of captured labels in the test region. It ultimately impacts the sensitivity and accuracy of the assay. By locally regulating the lateral flow speed, one can optimize the conditions for analyte detection, ensuring the necessary reaction durations. This fine-tuning of reaction kinetics directly influences the intensity of color change at the detection site, which is crucial for accurate target analyte quantification. Moreover, controlling the lateral flow speed allows for better control over the concentration of reactants along the flow path, leading to improved reproducibility and reliability of results. Overall, this strategy enhances the performance of LFAs by maximizing sensitivity, specificity, and consistency in analyte detection [22].

In this work, a straightforward approach for enhancing the reaction time and constraining the analyte flow in NC membrane by a varying number of parallel vertical µ-channels imposed employing femtosecond laser µ-machining was proposed. It was demonstrated that µ-channel density, width, and length can controllably constrain the analyte flow. Local delay of the flow by 950% resulted in boosting the immunoreaction rate between antigens and antibodies and increasing the number of captured labels. The colorimetric test line intensity readout of the SARS-CoV-2 analyte improved by 40% at defined µ-channel geometries.

## 2. Results and Discussion

### 2.1 Ablation Threshold of the NC Membranes

A porous NC membrane holds inherent surface inhomogeneities in the micron range and that imposes some challenges for laser ablation at certain resolution. For making precise microchannels in NC surface without reaching the polyvinyl chloride backing card it was necessary to determine the ablation threshold. It was obtained from a dedicated experiment where the second harmonic of Yb:KGW femtosecond laser (**Figure 1 a**) pulse energy was varied from 12.5 µJ to 57.5 µJ and the number of pulses was varied from 10 to 100.

**Figure 1.**
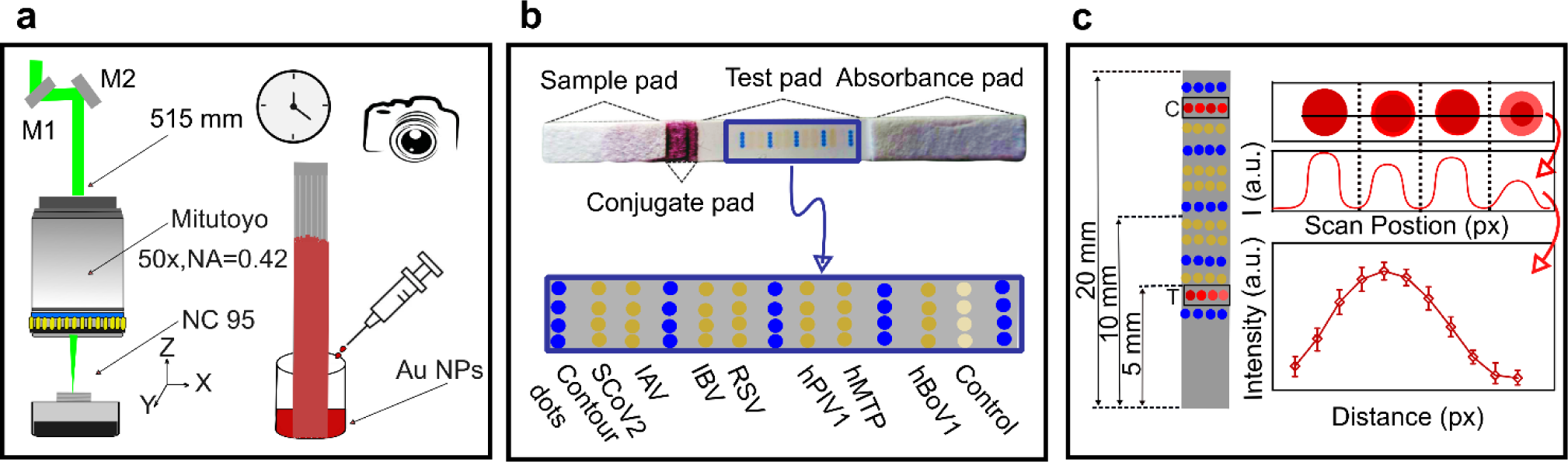
Schematic illustration of the experimental approach. (**a**) Laser µ-machining process used for structuring NC membrane employing the second harmonic beam delivered by mirrors. Wicking experiment setup for determining the µ-channel influence on the capillary flow rate. (**b**) Microarray LFA prototype test with antibody targets camera image with more detailed schematics below. (**c**) Automated sensitivity and signal enhancement analysis process from color images of test (“T”) and control (“C”) dotted lines averaging the intensity of the four dots.

By analyzing the relationship between the laser fluence (*F*) and SEM-measured hole diameter squared (*D*^2^) the laser beam spot radius *w*_0_ of 1.2 µm was deduced. Later on, from the intercepts the *F*_th_ for 10 and 100 puses was obtained 0.170 J/cm^2^ and 0.089 J/cm^2^, respectively (**Figure 2 a**). The ablated crater diameter ended up being bigger than the diffraction-limited spot size because the material is porous. Similar findings were observed in [26]. Hecht *et al.,* conducted a similar experiment where pulse energy of 0.882 μJ was required for the CN95 ablation with shorter pulse lenght femtosecond laser but it was obtained for the moving beam configuration with a f-Theta lens having 12 μm spot size [27]. Skordoulis, involved the use of a CO_2_ laser, with ablation thresholds reported at 3.0 and 4.1 J/cm². A moving beam configuration with 24 cm focal length optics was used in conjunction with an iris diaphragm to ablate the target [28].

**Figure 2.**
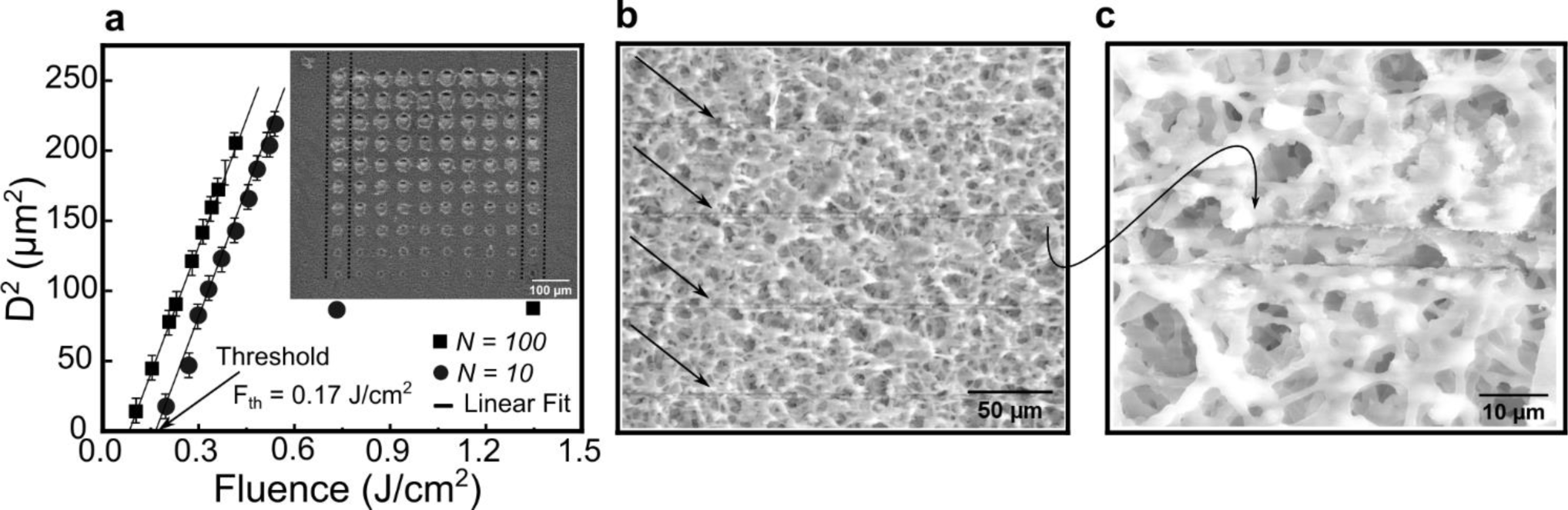
NC membrane ablation threshold determination. (**a**) The average squared crater diameter *vs.* applied laser fluence using *N*=10 and 100 pulses. The error bars represent one standard deviation from three diameter measurements. The inset depicts an fs-laser ablated membrane SEM micrograph with different energy and the number of pulses. (**b**) SEM micrograph of multiple laser µ-machined channels indicated by arrows. (**c**) Close-up SEM view of a single 5 µm width µ-channel.

By varying the laser pulse repetition rate and the scan speed two different conditions termed “Fast” and “Slow” ablation were analyzed in more detail (see **Table 3)** resulting in different ablation depth and line edge roughness as seen in **Figure 3**. It was obtained that under slow conditions the original membrane structure is intact and does not change the porosity. By using this more precise and controlled cutting method, we can avoid irregularities such as zigzags seen in the case of Fast cutting (Figure 3 bottom µ-channel). Slow ablation provides better-defined channels that reduce flow turbulence and resistance caused by uneven surfaces [27], despite longer µ-machining duration. Eventually, it might lead to better-controlled fluid flow. Also, it is advisable to perform an ablation close to the threshold conditions limiting the damage to the NC membrane.

**Figure 3.**
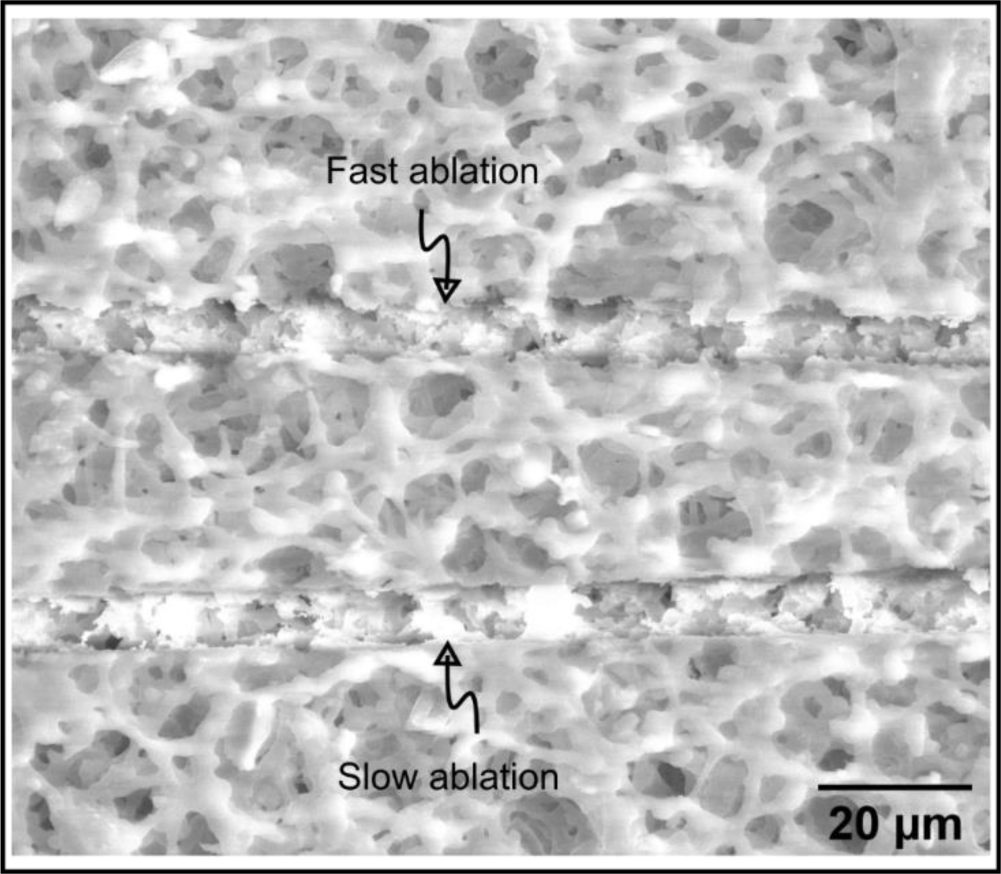
SEM micrograph of laser-ablated microchannels under two different conditions. The top µ-channel is abated by faster scanning than the bottom one as explaind in **Table 1**.

**Table 1.**
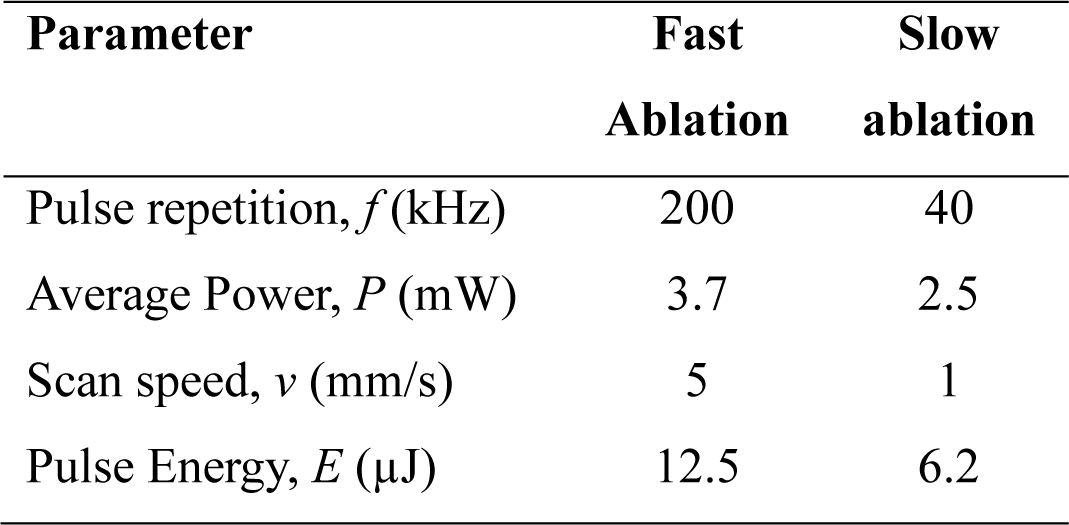
Typical laser parameters used for the ablation of NC membranes.

### 2.2. Characterization of Gold Nanoparticles

Aiming to mimic the LFA operation and wicking restriction in laser µ-structured NC membrane gold colloids were chemically synthesised. The synthesis of AuNPs was achieved using trisodium citrate as the reducing agent for Au NPs. The reduction reactions were conducted under carefully optimized conditions including controlled temperature and reaction time to facilitate the nucleation and growth of stable nanoparticles. The colloid had a characteristic pink color (**Figure 4 a** inset) explained by the LSPR-related Au NP extinction at 530 nm seen in the UV-Vis spectra (**Figure 4 a**) [29]. The particle shape was close to spherical and the average diameter was ca 50 nm (**Figure 4 b**).

**Figure 4.**
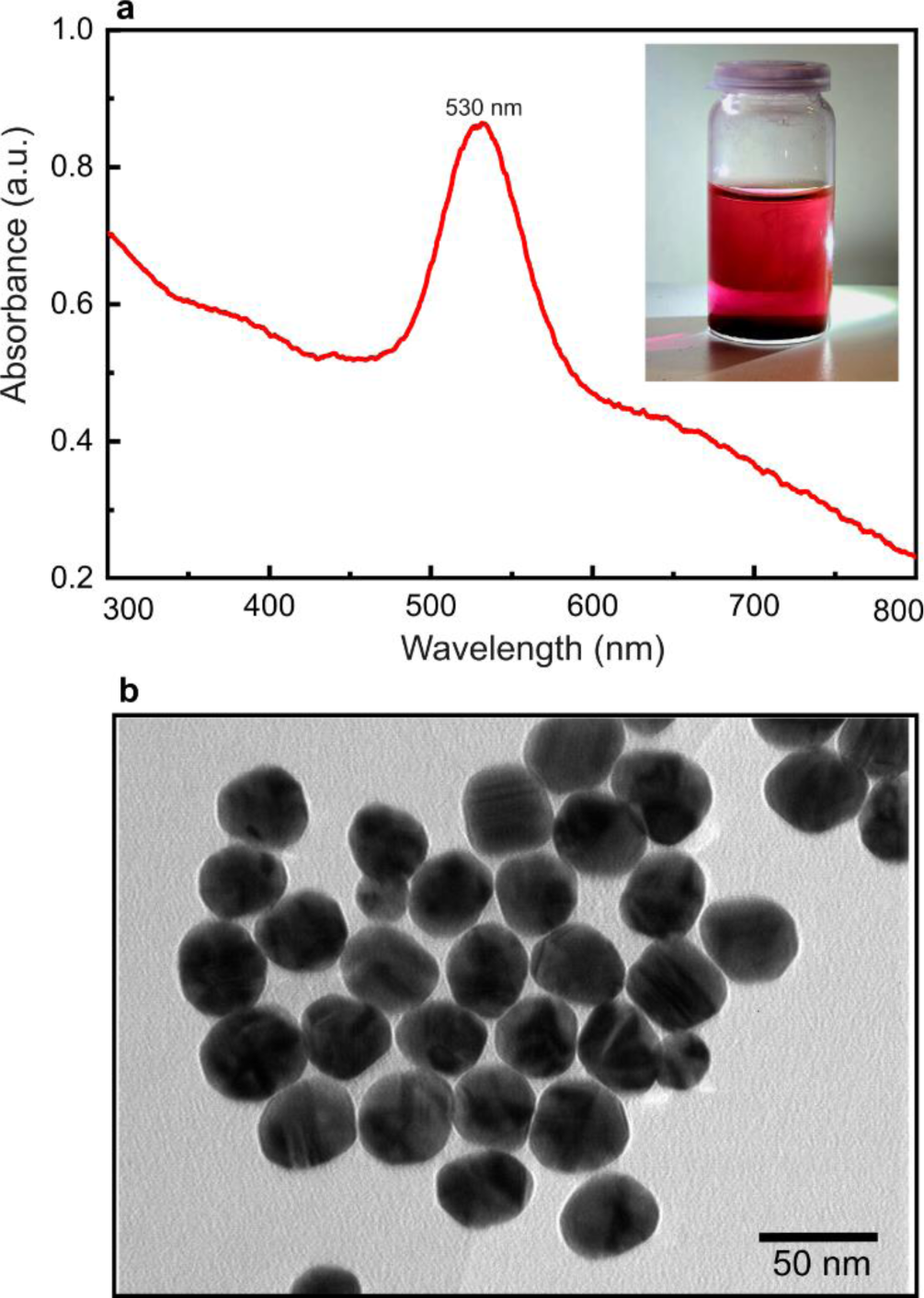
Characterization of gold NPs. (**A**) The UV-Vis absorbance spectra of AuNPs. The inset depicts the camera image of the gold NPs in the vial. (**b**) A TEM micrograph Au NPs.

### 2.3. µ-Channel Influence on the Wicking Flow

After selecting the “slow” laser ablation condition parameters (**Table 1**) the influence of the µ-channel width and length on the Au NP colloid vertical capillary flow (**Figure 1 a**) over a 20 mm length NC membrane (**Figure 1 c**) was investigated. The influence of the fs-laser µ-machined channel length and density is depicted in **Figure 5**.

**Figure 5.**
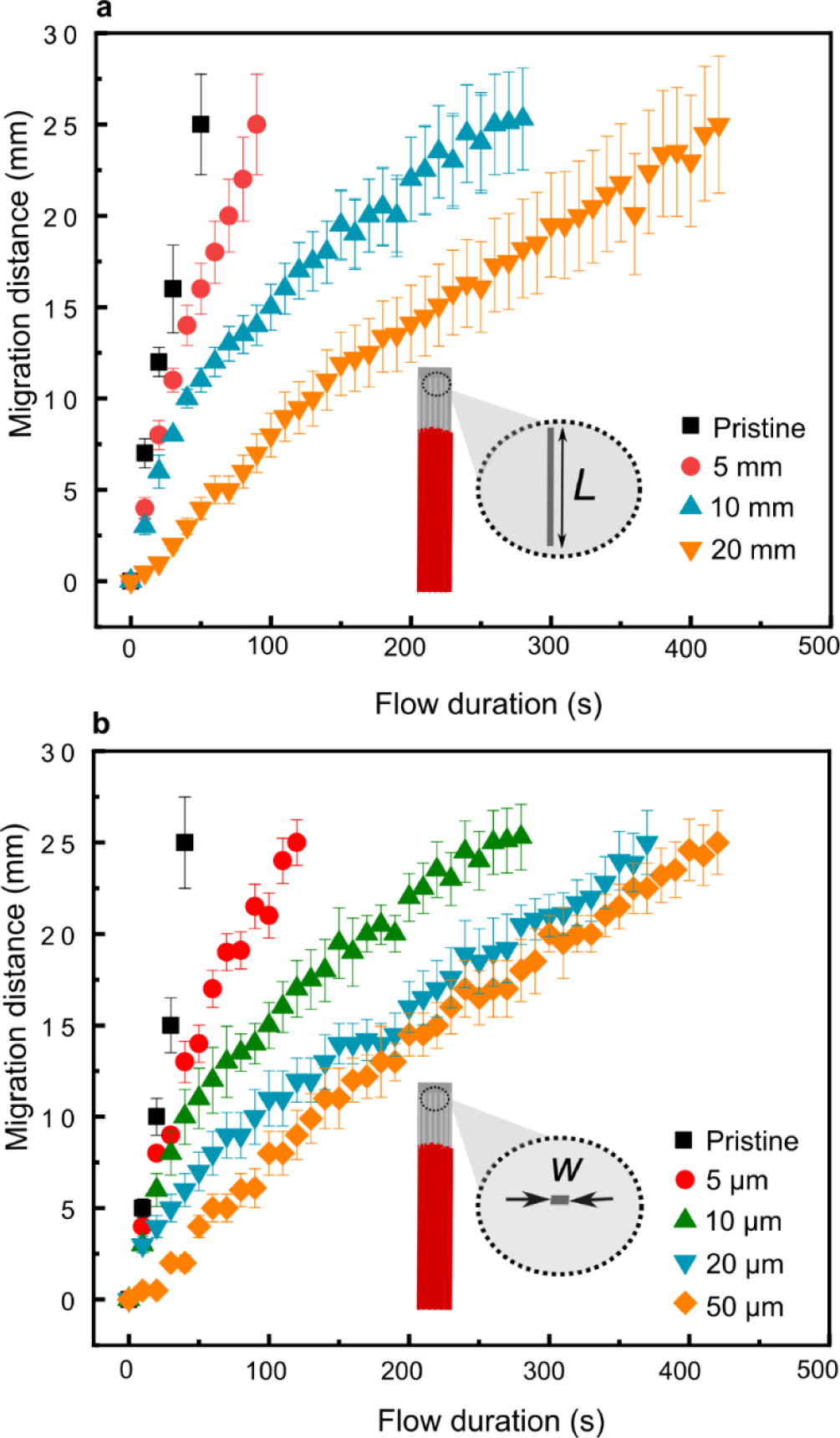
Migration distance as a function of time for pristine and laser µ-machined NC membranes. The migration distance of the Au NP colloid in NC membrane at different time moments depending on the length **(a)** and width **(b)** of laser ablated µ-channels. The µ-channel density in both cases fills 4 mm of the NC strip and the number of µ-channel depends on their width (gap and spacing are the same). The width in (**a**) is 50 μm and the length in (**b**) is 10 mm.

The nominal capillary vertical flow duration across a 20 mm long membrane was 40 s. Supplementary **Movie S1** and **Movie S2** depict camera videos of two typical wicking experiments of pristine and laser-treated NC membranes respectively. After the laser treatment, the wicking across the same length took up to 420 s as seen in **Figure 5**. Our study revealed that the laser treatment led to a substantial increase in the nominal flow rate, with the wicking process across the same length taking up to 380 s. It was estimated that the interaction time rose by 950% in comparison to the pristine strip when the µ-channel width was 50 µm and length was 20 mm. Shorter μ-channels of 10 mm and 5 mm length exhibited an increase of at least 600% and 125%, respectively. It was confirmed that varying the width of channels also has a noticeable impact (**Figure 5 b**) where μ-channels ranging from 5 to 50 µm exhibited a consistent reduction in wicking velocity, decreasing flow speed from 0.13 mm/s to 0.06 mm/s, respectively, when compared to 0.20 mm/s velocity in the unmodified NC.

This represents a significant deviation from previous methodologies, such as the approach by Fei Li *et al.*, where the electrospun polycaprolactone (PCL) on NC membrane resulted in a considerably shorter delay of flow, achieving 28±2% delay for travel a 3.0 cm distance [23].

In comparison to the work by Joong Ho Shin *et al.*, our study resulted in a similar delay compared to the NC membrane under compressive 23.54 MPa pressure which reduced the flow rate and led to a substantially longer time of approximately 11 more minutes for the diluted serum to flow through [30]. Xiao Feng Xu *et al.* [31], achieved delays of 25%, 75%, and 100% by integrating the sponge shunt after the conjugation pad into LFA and varying the thickness, length, and hydrophobicity of the sponge. Arben *et al.* [32], designed different wax barriers of different widths of 0.01 mm, 0.03 mm, and 0.05 mm and achieved 4 min, 7 min, and 12 min flow delay for lateral flow over 60 mm×3 mm membrane. Zhiqing Xiao *et al.* [33] developed an easy method to adjust the capillary flow rate on LFA substrates by using tape to cover the surface of substrates and achieved an average flow rate decrease to 61% of the original flow rate on synthetic paper.

### 2.4. µ-Channel Influence on the SARS-CoV-2 Detection Sensitivity

The experience gained on delaying flows in the laser-processed NC membrane was tested on the multiple respiratory diseases detecting LFA prototype based on the same type NC membrane using SARS-CoV-2 analyte. The LFA contained all the typical LFA components including sample pad, conjugate, and absorbent pads, test, and control lines as shown in **Figure 1 b**. The blue dotted lines in the diagnostic tests are for coordination and easiness of the readout process.

Aiming to distinguish the influence on the colorimetric readout of the laser μ-machined channels, the differently laser-treated LFAs were compared with the pristine LFA sample. The biofunctionalized active area of the test was laser treated with different lengths and density µ-channels employing the identical conditions met in **Section 2.3**.

After the SARS-CoV-2 analyte was applied to the pristine and laser-structured LFAs their color image was scanned and analyzed automatically using a custom image analysis code as explained in (**Figure 1 c**). The results of the Test and Control line relative intensities of differently laser-treated LFAs are summarized in **Figure 6**. The analyzed images of the Test and Control lines are depicted on the top rows in **Figure 6** together with the peak signal values corresponding to the brightest signal in the middle of the red dots used for colorimetric recognition of the specific binding event below. The averaged signal variation over the four dots widths is depicted in **Figure 7**.

**Figure 6.**
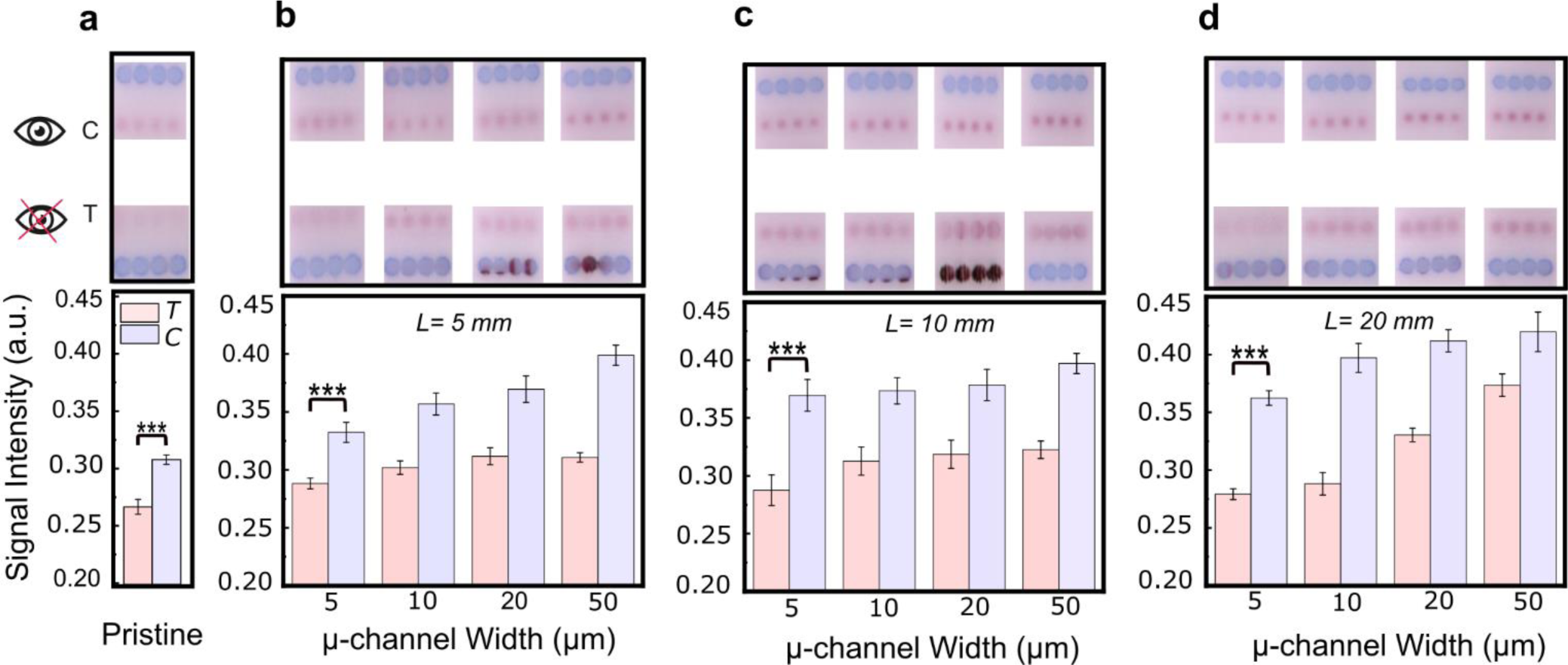
Maximum response of relative intensity of test (“T”) and control (“C”) lines as discerned from the camera image analysis after applying SARS-CoV-2 analyte. The signal intensity value is a grayscale amplitude averaged along the four dots. The error bars indicate the standard deviations of the gray-scale values obtained from four individual dots. The reference LFA without laser modification (**a**) and with laser µ-machined microchannel of different lengths 5 mm (**b**), 10 mm (**c**), and 20 mm (**d**).

**Figure 7.**
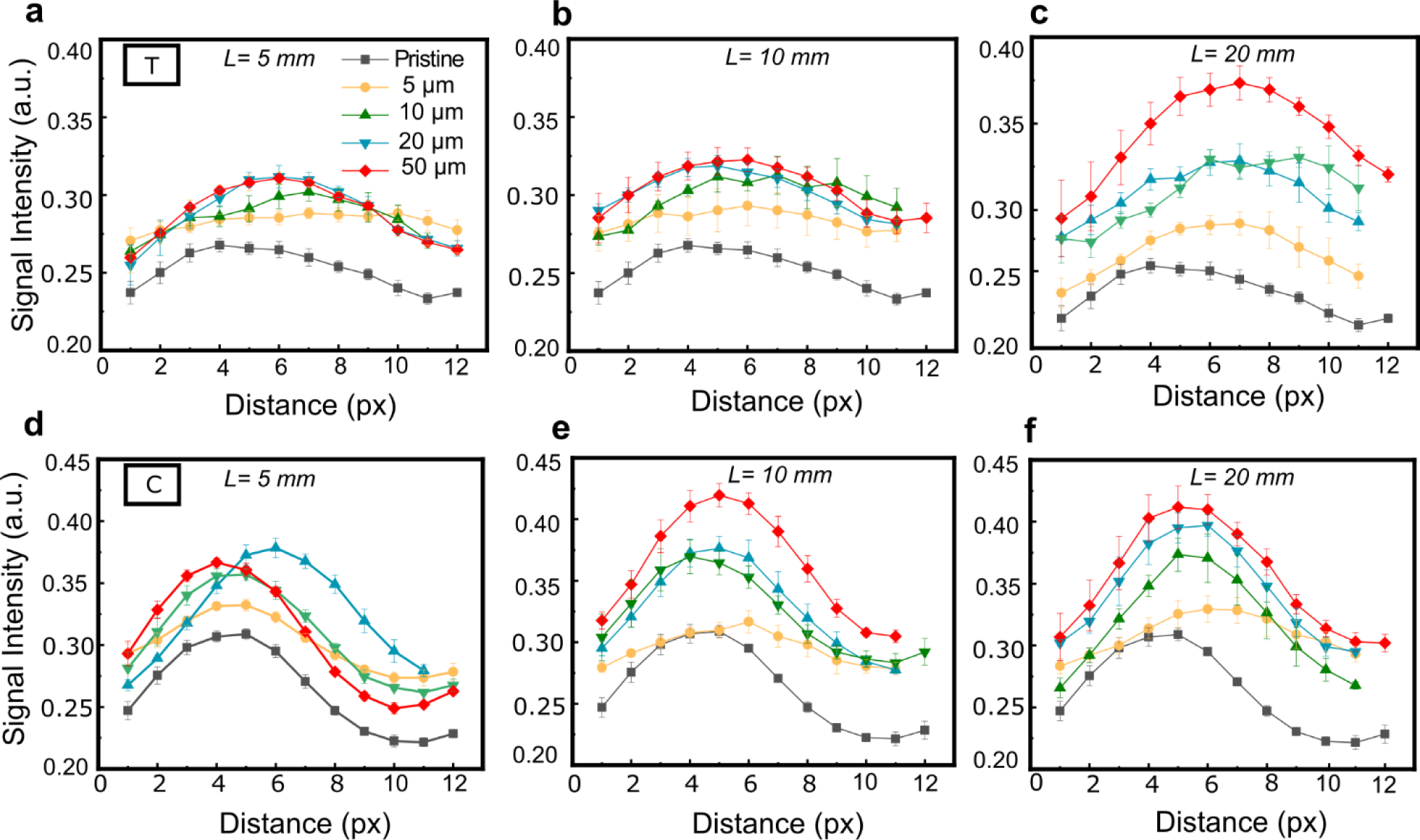
Pristine and differently laser μ-machined LFA “T” (**a**, **b**, **c**) and “C” (**d**, **e**, **f**) line image analysis results after applying SARS-CoV-2 analyte. Mean signal intensity was estimated from the automated analysis of the gray-scale value intensities measured along the four dots. The with of the channels is indicated in the legend and the length of the μ-channels was 5 mm (**a**, **d**), 10 mm (**b**, **e**), and 20 mm (**c**, **f**).

Low laser fluence and sparse µ-channels did not affect biofunctionalized areas during the engraving process of the NC membrane.

The SARS-CoV-2 test line intensity increased by 16% for 5 mm length and 50 µm width treated LFA (**Figure 6 b, Figure S3**) compared to the pristine test (**Figure 6 a**). When channel length was 10 mm, the intensity increased by 21% (**Figure 6 c, Figure S3**). The biggest signal was registered for a full strip length modified LFA (20 mm) where the test line signal increased by 40% for the case of widest μ-channels (**Figure 6 d, Figure S3**).

In the case of pristine LFA, the difference between the Test and Control lines was minimal because quite a big analyte concentration was applied. The microstructured LFA Test and Control line signal compared to the pristine sample increased in all investigated cases. It was observed that the signal was increasing with the increasing width of the µ-channels despite the overall length of the imposed modifications (**Figures 6**, **Figure 7**). The SARS-CoV-2 test line intensity increased by 16% for 5 mm length and 50 µm width treated LFA (**Figure 6 b**) compared to the pristine test (**Figure 6 a**). When channel length was 10 mm, the intensity increased by 16% (**Figure 6 c**). The biggest signal was registered for a full strip length modified LFA (20 mm) where the test line signal increased by 40% for the case of widest μ-channels (**Figure 6 d**).

Low laser fluence and sparse µ-channels did not affect biofunctionalized areas during the engraving process of the NC membrane. Still, slightly higher intensities over the test line were formed for 5 mm μLFA due to the narrow flow paths and slight local decrease of flow velocity (*v*), as seen in (**Figure 7 b)**. It can be observed that the binding of the AuNPs is more uniform in the case of 50 μm. In the case of 5 μm channel width multiplex LFA has not shown a noticeable difference in the brightness of the test line, as shown in (**Figure 7 a, b, c)**. In addition, the brightness of the test lines was higher when the length of channels increased as shown in (**Figure 6 d, Figure 7 c, f)**.

The results confirmed our initial hypothesis that µ-channel can constrain the Au NP flow velocity (*v*) through NC membrane flow before reaching the test region and increase the time of reaction (*T*). The laser modification increased the ratio of diffusion time to convection time of Au NPs (*Pe,* see **Table S1**) and decreased the ratio of the refraction flux to diffusion flux (*Da*), which is directly proportional to the reaction rate constant *k*^′^_*on*_ = *n* · *kon* for antibody-labeled Au NPs. Where *n* is an effective number of antibodies per Au NPs with calculated *Pe* >>1 (convection-dominated diffusion) and *Da*<<1 (diffusion-dominated reaction), thus the reaction was most likely a rate-limiting step to improve Au NP capture. While *k*_*on*_is the forward rate constant for a single antibody-antigen interaction in the NC membrane. The results suggest that significantly lowering and constraining the Au NP flow rate the prolonged reaction time leads to more expressed binding of AuNPs over the Test line.

## 3. Experimental / Materials and Methods

### 3.1 Used Materials

The nitrocellulose membranes used for laser treatment influence on wicking experiments in this study were NC 95 (Carl ROTH, Gmbh&CO) with an average pore size of 5 µm. The same NC membranes were used in the prototype multiplex respiratory viral diagnostic tests “MultiRespiTest” developed by UAB “Sanpharm” (Lithuania). The prototype diagnostic tests could detect specific binding with SARS-CoV-2 (SCoV2), Influenza B virus (IBV), Human Bocavirus 1 (hBoV1), RSV (Respiratory Syncytial Virus), hPIV1 (Human Parainfluenza Virus 1), hPIV3 (Human Parainfluenza Virus 3), and hMTP (Human Metapneumovirus). Only SARS-CoV-2 analyte was used in this work.

### 3.2 Methods

#### 3.2.1 Modification of NC Membrane by Laser µ-Machining

The femtosecond laser ablation tests and µ-channels in NC membranes on polyvinyl chloride backing card and prototype LFAs (MultiRespiTest, Sanpharm) were imposed employing a second harmonic of the 290 fs pulse length Yb:KGW femtosecond laser Pharos (Light Conversion, Lithuania) using an XYZ translation stage based workstation FemtoLab (Altechna R&D, Lithuania) which was controlled by the SCA µ-machining software (Altechna R&D, Lithuania) (**Figure 1 a**). The 515 nm wavelength light was focused with the 0.42 NA, 50x Plan Apo NIR objective (Mitutoyo, Japan). More details on the used laser system can be found elsewhere [34].

The ablation threshold of the NC membrane was determined by percussion-ablating an array of holes varying pulse energy from 12.5 µJ to 52.5 µJ with an increment of 10% and changing the number of pulses pulses from 10 to 100 with 10 pulse increments.

The different lengths µ-channels of 5 mm, 10 mm, and 20 mm were ablated in NC membranes, and LFAs as overlapping lines with 0.001 mm spacing were imposed under two characteristic ablation conditions that were analysed in more detail as summarized in **Table 1**. The number of µ-channels per 5 mm NC membranes or LFAs depended on the width of the single µ-channel keeping the gap of the same size. Microchannel widths of 5 µm, 10 µm, 20 µm, and 50 µm were investigated.

Laser-processed NC membranes were analyzed with an environmental field emission gun scanning electron microscope (SEM) Quanta 200 FEG (FEI, Netherlands) operated in the low vacuum mode.

#### 3.2.2 Preparation of AuNPs

Colloidal gold nanoparticles for wicking experiments were chemically synthesized according to the route described in [29]. In short, the synthesis consists of two steps starting from seed synthesis and the second step of growth of nanoparticles. Seeds were synthesized by heating 150 ml of 2.2 mM sodium citrate solution for 15 min under vigorous stirring. After the solution started to boil, 1 ml of HAuCl_4_ (25 mM) was injected. The color of the solution changed to soft pink in 10 min. Immediately after the seed synthesis, the reaction was cooled to 90°C, and 1 ml of HAuCl_4_ (25 mM) was injected. The solution was left for 30 minutes stirring heating at 90°C. This procedure was repeated twice. After that, 55 ml of sample was extracted and to the same vessel, 53 ml of water and 2 ml of 60 mM sodium citrate was added. Chemicals were acquired from Sigma Aldrich. This solution was then used as a seed solution, and the process was repeated. The absorbance of the colloid was inspected using UV-Vis-NIR spectrophotometer Avaspec-2048 and combined Deuterium/Halogen lamp light source AvaLight-DHc (Avantes, Netherlands). For the wicking, the colloid of 1.5 optical density (OD) was used as produced.

#### 3.2.3 Wicking Experiments

The vertical wicking speed of the pristine and laser-treated NC membrane strips of 5 mm width and 20 mm length was inspected by immersing in the 0.5 ml vial (Eppendorf) with 100 µl of the Au NP colloid. The process was recorded by a smartphone camera. The red color of the Au NP colloid improved the flow visualization and mimicked the actual LFA experiment. The measurement setup is depicted in **Figure 1 a**. The recorded videos were analyzed every 10 seconds to measure the solution front position on the NC membrane.

#### 3.2.4 Automated Image Analysis

The SARS-CoV-2 sample (10 ng/µL of recombinant SARS-CoV-2 nucleocapsid protein (UAB Baltymas, Lithuania) in 100 µL sample buffer) was applied on pristine and differently laser-treated LFA prototype tests that were then scanned with a flatbed scanner Perfection V370 Photo (Epson) for further analysis. The doted color test lines were carefully chosen in the images, and the average intensity values were measured by a dedicated code implemented in MATLAB (MathWorks, CA, USA). The key steps of the region of interest selection and image pretreatment are showcased in **Figure S4**. The signal values were evaluated by removing the background noise by calculating the average intensity value for each doted strip. The pixel intensity across the line, originally stored as a full-color RGB, was converted to grayscale. The resulting intensity values were inverted to address a darker color to a higher intensity value. A slicing location of the affected region was selected manually and a single pixel-wide line passing through all 4 dots of the Test line and Control lines was extracted for further analysis. The originally dotted line was then divided into 4 equal-length segments of 12 pixels. The segments were averaged and the standard error of the mean was computed for each pixel value, including the maximum value of the average. The Origin 2019b (OriginLab) software was used for the result statistical analysis.

## 4. Conclusions

We have proposed an innovative approach to improve the signal sensitivity of colorimetric readout lateral flow assays (LFAs) based on nitrocellulose membranes. It was achieved by imposing microchannels in NC membranes employing femtosecond laser cold ablation. The microchannels delayed the flow rate on the strip thereby increasing reaction time between the analyte and the labeled antibody leading to a more effective formation of the immunocomplex observed as a brighter colorimetric signal. The 50 µm widths µ-channel demonstrated an increase in flow rate delay of 125% and 600% when the channel length was extending over 5 mm and 10 mm, respectively. Moreover, this value reached 950% when the channels extended over all 20 mm lengths of the NC membranes. As a result of this strategy, the sensitivity of the prototype LFA test was increased by 40% for the SARS-CoV-2 analyte compared with the untreated LFA. The proposed method can be extended for further exploring the limit of detection and analytical sensitivity of point-of-care diagnostic devices.

## Contributions

Gazy Khatmi (ORCID id: 0000-0001-6989-9286) – Investigation, Formal analysis, Visualization, Writing – original draft preparation, Writing – review, and editing.

Tomas Klinavičius (0000-0002-7925-1691) – Methodology, Writing – original draft preparation.

Martynas Simanavičius (0000-0003-2996-229X) – Conceptualization, Methodology, Investigation, Formal analysis, Writing – review, and editing.

Laimis Silimavičius (0009-0005-8368-8350) – Conceptualization, Methodology, Writing – review, and editing.

Asta Tamulevičienė (0000-0003-4152-1382) – Investigation, Writing – original draft preparation.

Agnė Rimkutė (0009-0003-7022-3701) – Methodology, Investigation, Formal analysis

Indrė Kučinskaitė-Kodzė (0000-0002-1761-0089) – Methodology, Investigation, Formal analysis Gintautas Gylys – Conceptualization, Methodology, Project administration.

Tomas Tamulevičius (0000-0003-3879-2253) – Conceptualization, Visualization, Methodology, Resources, Funding acquisition, Project administration, Writing – original draft preparation, Writing – review, and editing.

## Acknowledgments

This work was supported by the project No. BIOTECH-02-014 under the measure “Development of Biotechnology Industry in Lithuania”.

## Ethical Approval

This research did not involve human or animal samples.

## Availability of data and materials

The datasets generated during and/or analysed during the current study are available from the corresponding authors on reasonable request.

## Supplementary Information

It contains Figure S1 depicting the LFA working principle, equations (S1)-(S5) describing the mass transport theory, Table S1 summarizing numbers defining the transport and reaction kinetics, Table S2 providing relevant parameter values for LFAs, Figure S3 depicting analysis of the relevant signal intensities, and Figure S4 explaining the automated image analysis sequence. Supplementary movies S1 and S2 showcase the differences in wicking of pristine and laser-treated nitrocellulose membranes.

