## Supplemental for "Lateral Flow Assay Sensitivity and Signal Enhancement via Laser µ-Machined Constrains in Nitrocellulose Membrane"

### 1. Working Principles of the Lateral Flow Assay

In their fundamental design Lateral flow assays (LFAs) are characterized by a conjugate comprising a signal particle and a binding molecule dried down on a porous pad that is placed on top of a membrane (**Figure S1**). Several drops of sample containing the analyte are placed on top or in front of the conjugate, with the analyte reacting with the conjugate as the resulting complex flows laterally along the membrane. A capture dots / line comprising a second binding reagent to the analyte then captures the complex, resulting in a visible line.

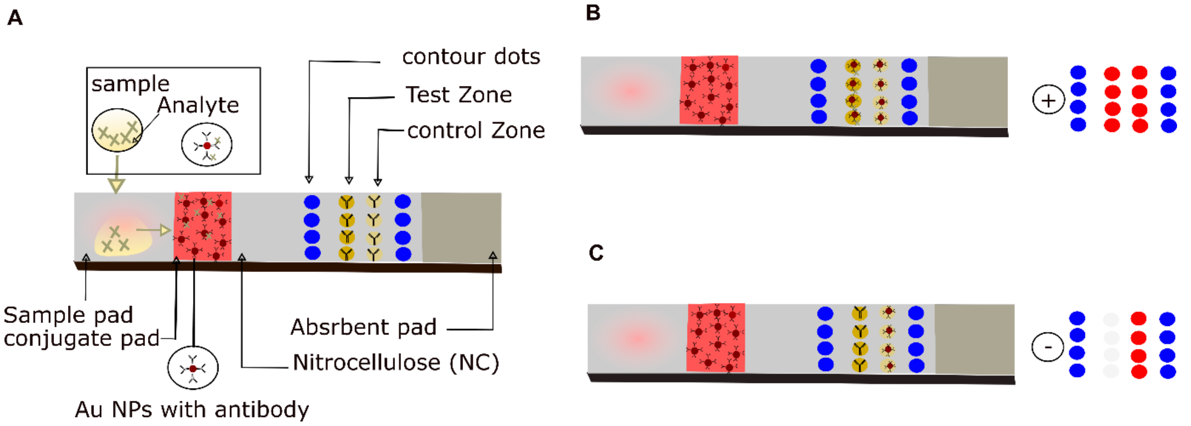


**Figure S1.** A schematic depicting the placement of reagents and porous membranes in a typical lateral flow test. **A** sample addition. **B** describes the antibody-antigen-signal particle sandwich

complex present at the test line for a positive sample, whereas **C** shows the results for a negative sample.

### 2. Theoretical Description of the Lateral Flow in the Membranes

The efficacy of LFAs hinges on several critical elements. One of the main factors is the precise timing of the reactions between the binding reagents and the analyte while minimizing nonspecific binding of analyte reagents, as well as maximizing the visual signal resulting from the aggregation of analyte–conjugate complexes on the test line [1].

Mass transport and reaction theory are fundamental in the development of lateral flow assays (LFAs) mechanistic theories. By understanding the movement and reaction of molecules within the assay structure, these theories can enhance the design and functionality of the assay [2].

Mass transport theory primarily investigates the movement of reactants (such as analytes and conjugates) over the nitrocellulose membrane of the LFA. This movement is regulated by diffusion, capillary action, and potentially, convection. Reaction theory, however, concerns itself with the interactions between these reactants and particular receptors on the test line, specifically focusing on the rate at which they bind together and the durability of the complexes they produce.

#### 2.1 Key Aspects in Mass Transport Theory

##### 2.1.1 Advection, Diffusion, and Dispersion of Reactants within LFA

Transport of reactants through nitrocellulose membranes to test line receptors is essential. In order for LFA to function properly this movement of reactants in an LFA from the location of sample input to the test line occurs through advection, diffusion, and kinematic dispersion.

The diffusion of reactants in lateral flow assays (LFAs) is primarily governed by Brownian motion, where molecules move randomly but tend to drift from areas of higher concentration to lower concentration, following concentration gradients.

The porous structure of nitrocellulose membranes significantly impacts the diffusion behavior of reactants within the assay. The effective diffusion coefficient $D_{eff}$ for reactant n can be defined as [2, 3]

$$D_{eff,n} = \frac{\varepsilon}{\tau} D_n \qquad \text{(S1)}$$

where $\boldsymbol{\varepsilon}$ is the porosity of the material, $\tau$ is the material tortuosity, and $D_c$ is the free solution diffusion coefficient for reactant $n$.

The advection-dispersion equation is used to describe the total transport of reactants through the LFA by combining the effects of advection and dispersion.

$$\varepsilon\frac{dC_n}{dt} = D_{di}\, \nabla^2 C_n - \mathrm{V}\nabla \mathrm{C_n} \qquad \text{(S2)}$$

where $C_n$ is the concentration of reactant $n$, $V$ is the velocity vector, and $D_{di}$ is the dispersion coefficient [4].

#### 2.1.2 Analyte Binding to Antibody-Coated Au NPs

When an analyte is added to the sample pad, antigen-signal particle complexes dominate the test line signal, especially in samples with low antigen concentrations. The liquid-phase reaction between an antigen and an antibody-coated nanoparticle represents reversible ligand binding to cell surface receptors. The theory of ligand binding to cell surface receptors is explained by Berg & Purcell [5].

$$k_{on,Particle} = \frac{4\,\Pi\, a\, D_{ef}\, N k_{on}}{N k_{on} + 4 \Pi a D_{ef}} \qquad \text{(S3)}$$

$$K_{off,particle} = k_{off}\left(\frac{1-4\Pi a D_{ef} N k_{on}}{N k_{on} + 4\Pi a D_{ef}}\right) \qquad \text{(S4)}$$

where $N$ is the number of receptors on the cell surface, $a$ is the cell radius, $k_{on}$ is the association rate for complex formation, and $k_{off}$ is the dissociation rate constant.

At the test line, a number of events can result in the formation of signal particle complexes. The two main routes are, binding of antigen-nanoparticle complexes to available test line receptors (antibodies), or by a sequential binding reaction, where an antigen binds to the test line receptor, and a nanoparticle binds to the antigen-receptor complex. This is described by:

$$\boldsymbol{\varepsilon}\frac{dC_n}{dt} = D_{di}\nabla^2 C_n - \mathrm{v}\nabla C_n + R_n \qquad \text{(S5)}$$

$R_n$ is the local rate of change of antigen due to the reaction.

#### 2.2. Reaction Regimes Within LFA

The kinetics of transport and immunological reaction are characterized by the Péclet number ($Pe$) and the Damköhler number ($Da$), respectively.[6] Where The Péclet number is a dimensionless number that characterizes the relative importance of advective (or convective) transport to diffusive transport. The Damköhler number is a dimensionless number that compares the chemical reaction rate to the rate of transport processes **(Table S1** and **Table S2).**

**Table S1** Numbers defining the transport and reaction kinetics in the LFAs

| Dimensionless number | Equation | Description |
|---|---|---|
| Péclet number ($Pe$) | $vS/D$ | Advective rate / diffusion rate |
| Damköhler number ($Da$) | $C_e\ K_{on}\ S/D$ | Reaction rate/diffusion rate |

**Table S2** Relevant parameter values for LFAs

| Parameter | Description | Typical values | Reference |
|---|---|---|---|
| $S$ (µm) | Nitrocellulose pore radius | 0.2 – 10 | [7] |
| $v$ (m/s) | Average fluid flow velocity | 0.5 – 3 ×$10^{-4}$ | [7, 8] |
| $D$ ($cm^2/s$) | Diffusivity (molecules conjugation labels) | $10^{-5}$ – $10^{-8}$ | [3, 8, 9] |
| $C_e$ (ng/ml) | The concentration of captured molecules in the test region. | 0.01 – 1.0 | [10, 11] |
| $K_{on}$ ($M^{-1}S^{-1}$) | Reaction rate constant | $10^4$ – $10^7$ | (13–15) |

The relative test line signal increase compared to laser untreated LFA is depicted in **Figure S3**

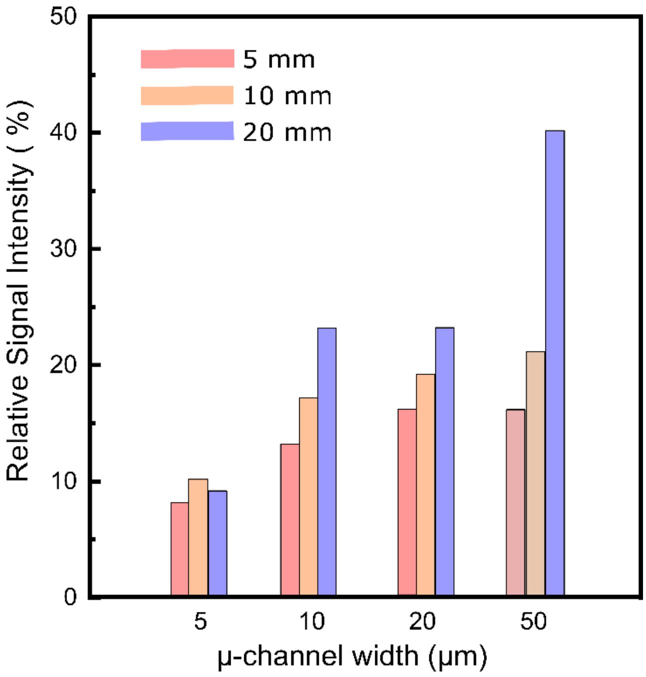


**Figure S3.** The relative SARS-CoV-2 analyte signal intensity increase of differently laser μ-machined LFA with 5 mm, 10 mm, and 20 mm length μ-channels with respect to pristine NC membrane.

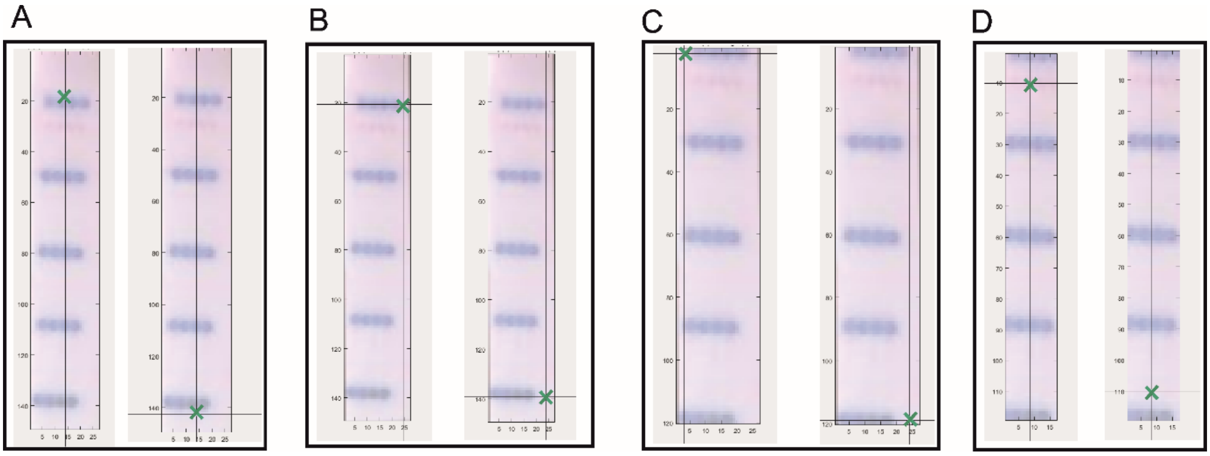


**Figure S4**. Image analysis steps. **A** Tilt/rotation compensation of vertical image from two selected dots, **B** Horizontal crop, **C** Vertical crop, and **D** region of interest identification, where top pink dotted line is Control line and bottom is SARS-CoV-2 test line. Green "x" indicates the mouse click positions.
